# The tRNA Gm18 Methyltransferase TARBP1 Promotes Hepatocellular Carcinoma Progression via Metabolic Reprogramming of Glutamine

**DOI:** 10.1101/2023.06.14.544902

**Authors:** Xiaoyan Shi, Yangyi Zhang, Yuci Wang, Jie Wang, Yang Gao, Ruiqi Wang, Liyong Wang, Minggang Xiong, Ningjing Ou, Qi Liu, Honghui Ma, Jiabin Cai, Hao Chen

**Author notes:** These authors contributed equally to this work (Xiaoyan Shi, Yangyi Zhang, Yuci Wang, Jie Wang).

## Abstract

Cancer cells rely on metabolic reprogramming to sustain the prodigious energetic requirements for rapid growth and proliferation. Glutamine metabolism is frequently dysregulated in cancers and is being exploited as a potential therapeutic target. In current study, we identified TARBP1 (TAR (HIV-1) RNA Binding Protein 1) as a novel driver gene critical for glutamine metabolic reprogramming in tumor through the CRISPRi/Cas9 screening. Our *in vivo* and *in vitro* assays demonstrated that TARBP1 is the methyltransferase of Guanosine 2’-O-methylation targeting position 18 (G18) of tRNA^Gln^ ^(TTG/CTG)^ and tRNA^Ser^ ^(TGA/GCT)^, and loss of Gm18 modification diminishes the stability of tRNAs. Therefore, TARBP1 is critical for maintaining efficient translation of mRNA, in particular the glutamine transportor-ASCT2 (also known as SCL1A5). Importantly, TARBP1 is frequently amplified and overexpressed in HCC, consequentially promotes the protein synthesis of ASCT2 and glutamine import to fuel the growth of cancer cell, which is associated with poor patient survival. Taken together, this study reveals the critical role of TARBP1 in HCC progression through glutamine metabolic reprogramming and provides a potential target for tumor therapy.

## INTRODUCTION

Hepatocellular carcinoma (HCC), accounting for 90% of primary liver cancer, is one of the most frequent malignancies in the world, with 0.9 million new cases and 0.8 million deaths globally in 2020^1^. However, HCC therapy remains a significant challenge, such as the high therapeutic resistance, and developing new strategies to treat HCC is a continued need^2^.

Tumor growth has prodigious energetic requirements. In order to fulfill fast cell growth and proliferation, cancer cells frequently rewire metabolic pathways to rapidly synthesize adenosine triphosphate (ATP) and increase biomass accumulation for biosynthesis^3^. Glucose and glutamine are the two major nutrients consumed by cancer cells. As over 90% glucose in proliferating cancer cells is preferentially metabolized to lactate (known as the Warburg effect), and limited pyruvate is transferred into the tricarboxylic acid cycle (TCA), glutamine can replenish metabolites to maintain mitochondrial function^4^. In addition, glutamine also serves as the major nitrogen and carbon donor for the biosynthesis of nucleic acids, proteins, and lipids^4, 5^. Thus, a growing body of evidence suggests that several cancers, including HCC, are addicted to glutamine, and targeting cancer glutamine metabolism has become a novel therapeutic approach^4–7^. However, many ways for blocking glutamine metabolism, including glutamine uptake inhibitors, glutaminase inhibitors, and glutamine antagonists, are generally limited^8^, suggesting the continued need to identify other or combinatorial strategies.

The key enzymes participating in glutamine metabolism are modulated by many oncogenes targeting gene expression regulation^4, 9, 10^. For instance, c-Myc, the third most frequently amplified oncogene, activates the transcription of glutamine transporters by binding to their promoter elements^9^ and upregulates GLS expression through transcriptionally repressing miR-23a/b that target the GLS 3’UTR^10^.

tRNA is essential for protein synthesis, which can recognize mRNA codons and bridge corresponding amino acids. It’s not surprising that dysregulated tRNA expression has been commonly observed in many cancer cells. However, the underlying mechanism is still largely unknown. Recent research advanced tRNA modifications are crucial for translation and cancer progression. For example, modifications that occur in wobble positions, such as ncm5U or mcm5s2U at position 34, can enhance base pairing to maintain the correct reading frame during codon-specific translation. Morever, m5C at position 34 in mitochondrial tRNA is required for protein translation of AUA codons. Additionally, other anticodon loop modifications increase the translation speed and fidelity of codon-specific genes. Modifications outside the anticodon loop also regulate tRNA biogenesis. G18m is one of most confusing tRNA modifications.

TAR (HIV-1) RNA Binding Protein 1 (TARBP1) protein with 1621 amino acids contains an SPOU domain and belongs to the SpoU-TrmD superfamily^1^^1,1^^2^ (Extended Data Fig. 1a). This methyltransferases family is mainly responsible for the modifications of tRNA and rRNA, including 2’-O-methylation (Nm), m1G, and m3U^1^^3^. Among them, Guanosine 2′-O-methylation at position 18 (Gm18) of tRNA is a highly conserved modification from bacteria to mammal, which is present in almost all T. thermophilus tRNA species, 13 E. coli tRNA species, while only two human tRNA species (Ser and Gln)^1^^4,1^^5^. The Gm18 of tRNA is catalyzed by TrmH in E. coli.^1^^6^, and Trm3 in S. cerevisiae^1^^7^. Recently, Freund. et al. reported that TARBP1, the human homolog of TrmH and Trm3, has the proposed 2 ′ -O-methyltransferase activity targeting tRNA at position 18 through RiboMethSeq analysis^1^^4^, but its substrate requirement and specificity, and the biological function of Gm18 remain largely undefined.

**Fig. 1:**
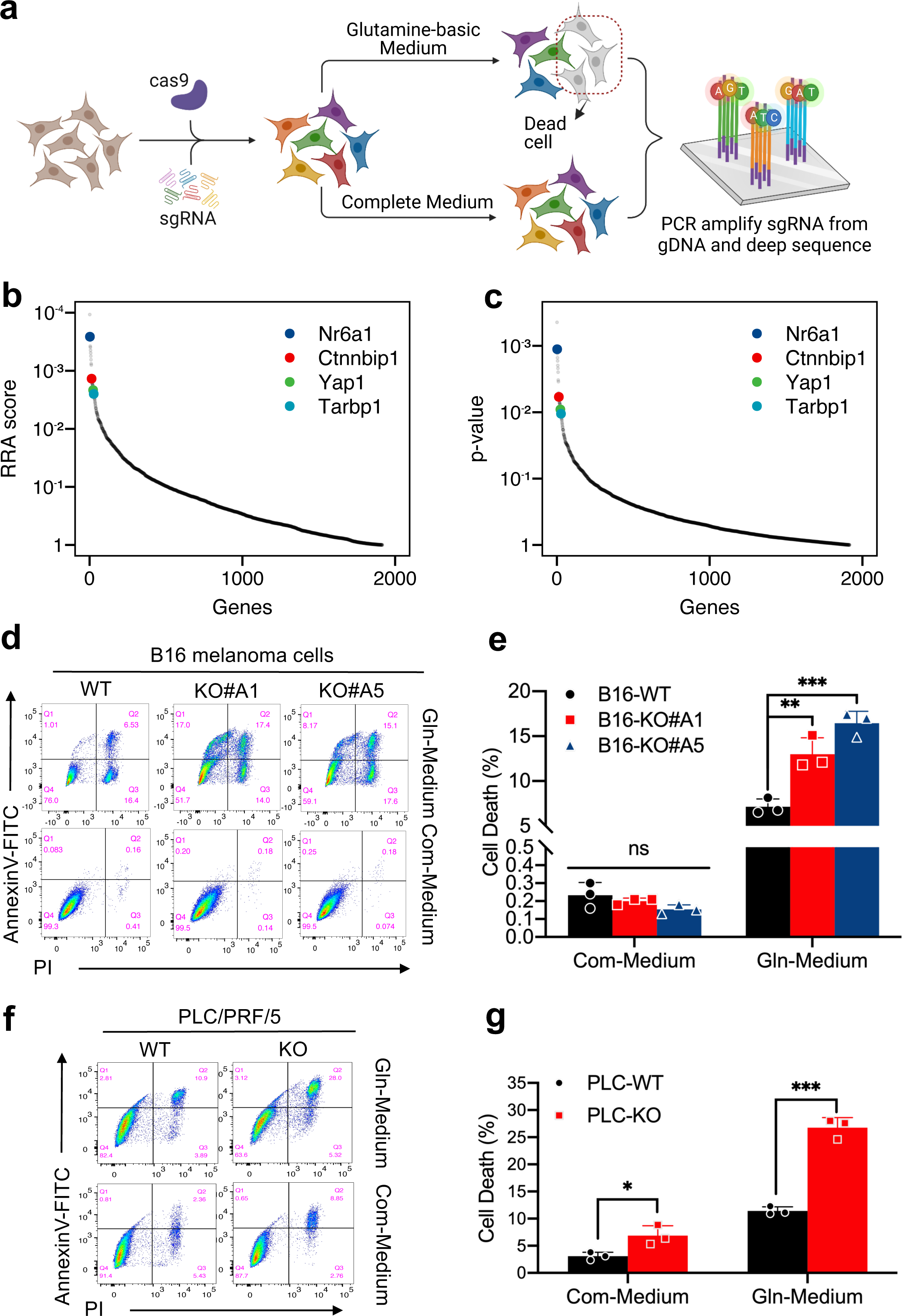
CRISPRi/Cas9 library screening identify TARBP1 as a novel regulator of cancer glutamine metabolism. a, Screening strategy. Cell populations with sgRNA library were cultured in complete medium (with glucose, pyruvate and glutamine) or glutamine-basic medium (only containing glutamine as the sole energy source) for 48 hours. Dead cells taken in glutamine - basic medium and cells taken in complete medium were subjected to next-generation sequencing to evaluate the abundance of sgRNAs. b, c, Rank-order plot of P-value (b) and gene scores (c) for the CRISPRi screen. d-g, Representative flow cytometry images (d,f) and quantification analysis (e,g) of the apoptotic B16 melanoma cells (d,e) and PLC/PRF/5 cells (f,g)culturing in glutamine-basic medium or complete medium for 48 hours. FITC: fluorescein isothiocyanate; PI: propidium iodide. *P<0.05, **P<0.01, ***P<0.001, ns: not significant. Student’s t-test. Data were presented as mean ± SD (n=3).

Here, using CRISPRi/Cas9 library designed for screening genes involved in expression regulation, we identified TARBP1 as a novel regulator for basic glutamine metabolism. Therefore, we reasoned that TARBP1 might play crucial roles in gene-specific translation through Gm18 modification of tRNA, thereby promoting glutamine metabolism reprogramming in cancer. To interrogate our hypothesis, we set out to characterize the 2 ′-O-methyltransferase activity of TARBP1 and investigate the molecular function of Gm18. Here, we demonstrated that TARBP1 mediates Gm18 methylation targeting tRNA^Gln^ ^(TTG/CTG)^ and tRNA^Ser^ ^(TGA/GCT)^ in vitro and in vivo, and Gm18 modification can enhance the stability of modified tRNAs. Depletion of TARBP1 caused global translation deficiency, affected the translation efficiency of ASCT2, and inhibited glutamine transport and tumorigenicity.

## RESULTS

### CRISPRi/Cas9 library screening identified TARBP1 as a novel regulator for glutamine metabolism

To systematically identify the novel genes involved in regulation of glutamine metabolism, a CRISPRi/Cas9 library with 10090 unique single-guide RNAs (sgRNAs) that target 1918 distinct protein-coding genes was used for performing the screening in a cell model. The B16-F10 melanoma cells were cultured in two conditions: complete medium (with glucose, pyruvate and glutamine) or glutamine-basic medium (only containing glutamine as the sole energy source) for 48 hours. We hypothesized that cells depleted of genes essential for glutamine metabolism will die in the glutamine-basic medium (without glucose and pyruvate). Hence, dead cells cultured in glutamine-basic medium and cells cultured in complete medium were collected to carry out next-generation sequencing for evaluating the abundance of sgRNAs (Fig. 1a and Extended Data Fig. 1c,d). Using the MAGeCK scores^1^^8^, we identified *Tarbp1* may be a novel regulator for basic glutamine metabolism (Fig. 1b,c). All sgRNAs targeting *Tarbp1* were remarkably increased in dead cells collected from glutamine-basic medium groups, comparing with control groups (Extended Data Fig. 1b). To confirm the results from our screening, we generated *Tarbp1* knockout clones in B16 melanoma cells (Extended Data Fig.1e) and cultured WT and KO cells in glutamine-basic or complete medium for 48 hours. Consistent with the high-throughput screening results, a significant increase in the number of apoptotic cells was observed in *Tarbp1* knockout cells cultured in a glutamine-basic medium but not in a complete medium (Fig.1d,e and Extended Data Fig. 1f). Then, we used PLC/PRF/5 - a human HCC cell line - to verify the results further. In line with results obtained in B16 cell line, we observed that the apoptotic cells from TARBP1 KO group were remarkably increased when culturing in the glutamine-basic medium (Fig.1f,g and Extended Data Fig. 1g), implying that TARBP1 is indeed involved in glutamine metabolism.

### TARBP1 is required in HCC progression

Given that tumor cells show increased consumption of glutamine and strong dependence on glutamine metabolism^4–7^, we asked whether TARBP1 will play a critical role in tumorigenesis. Subsequently, we analyzed TCGA database and found that TARBP1 is frequently amplified and overexpressed in various cancers (Fig. 2a,b). Noticeably, TARBP1 amplification is most common in breast cancer (∼ 10%), ovarian cancer(∼ 6.85%) and hepatobiliary cancer (∼ 6.72%). Importantly, TARBP1 expression is positively correlated with tumor stage and differentiation grade of HCC (Extended Data Fig. 2a,b)^1^^9^. Next, we sought to define the potential function of TARBP1 in tumorigenicity using liver cancer cell model. We knocked out TARBP1 in two different HCC cells lines - PLC/PRF/5 and HepG2 - employing CRISPR/Cas9 (Extended Data Fig. 2c) and validated the knockout efficiency of TARBP1 by Western blot (Fig. 2c, and Extended Data Fig. 2d). Consistently, TARBP1 deletion exerted a strong suppression of cell growth both in PLC/PRF/5 and HepG2 cells (Fig. 2d, and Extended Data Fig. 2e). Next, monoclonal formation assay and wound-healing assay were carried out, and we observed that depletion of TARBP1 decreases both colony-forming abilities and migratory abilities of HCC cell lines (Fig. 2e-h). Furthermore, cell cycle analysis revealed that the percentage of G0/G1 phase cells in TARBP1-KO cells increased, while the proportion of S phase cells decreased, suggesting that loss of TARBP1 significantly arrests cell cycle progression (Extended Data Fig. 2f,g). To further verify the tumorigenicity of TARBP1 *in vivo*, xenograft tumor-formation assay was performed. In line with results obtained in cell model, we discovered that TARBP1 depletion dramatically suppressed the tumor formation, suggesting that TARBP1 inhibition significantly impaired tumor growth *in vivo* (Fig. 2i-k). Collectively, our results provide evidences that TARBP1 plays a critical role in HCC progression.

**Fig. 2:**
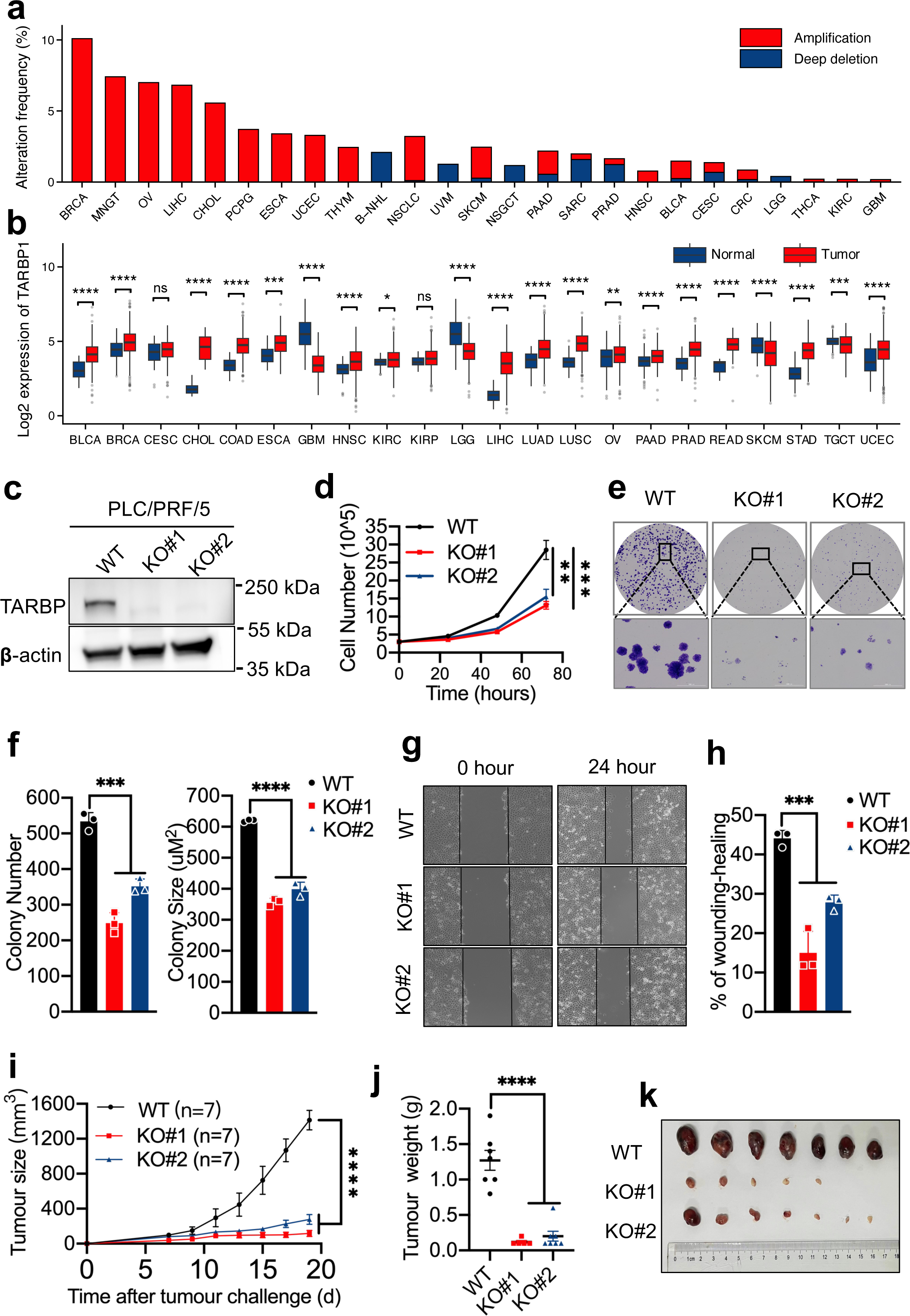
TARBP1 is necessary for cancer cell growth and oncogenicity. a, TARBP1 amplification in human cancers. b, Expression levels of TARBP1 in tumors and peri-tumor tissues in different cancer types. c, Western blot analysis confirming TARBP1 KO clones in PLC/PRF/5 HCC cell line by TARBP1 antibody. β-actin was used as a loading control. d, Cell proliferation analysis of WT and TARBP1 KO cells. e,f, Colony-formation assay of WT and TARBP1 KO cells. 1000 cells were seeded in 6-cm tissue culture dishes, representative pictures were taken (e), and the colony numbers and sizes (f) were measured on day 9. g, h, Wound-healing images of WT and TARBP1 KO cells. Representative pictures were taken (g), and the scratch area after healing (h) was measured at 24 hours. i-k, Overview of tumors in NUDE mice subcutaneously implanted with WT and TARBP1 KO cells. The tumor growth (i) and final tumor weights (j) were compared, and a picture of the final tumor were taken (k). *P<0.05, **P<0.01, ***P<0.001, ****P<0.0001, ns: not significant. Student’s t-test. b, Data were presented as mean ± SEM. d, f, h, Data represented as mean ± SD (n=3). i, j, Data were presented as mean ± SEM (n=7).

### TARBP1 is the *bona fide* Gm18 methyltransferase targeting tRNA^Gln^ ^(TTG/CTG)^ and tRNA^Ser^ ^(TGA/GCT)^ both *in vitro* and *in vivo*

TARBP1 contains a typical SpoU_methylase domain (Extended Data Fig. 1a), which is conserved from bacteria to mammals^1^^1,1^^2,1^^6^. Previous studies demonstrated that this SpoU_methylase doamin is likely responsible for the Gm18 modifications of tRNA in vaious species (Extended Data Fig. 3a)^13^. However, a comprehensive understanding concerning its enzymatic activity and molecular functions is still largely unknown, especially in vertebrates. Therefore, we aimed to investigate the roles of TARBP1’s enzymatic activity and TARBP1-mediated tRNA Gm18 modification in the oncogenic effects of TARBP1. Our results showed that decreased proliferation caused by TARBP1 deficiency can be partially rescued by expressing wild-type TARBP1 in KO cells, whereas expression of the catalytically inactive mutant (Mut) TARBP1 doesn’t have this effect (Fig. 3a,b). To verify the 2’-O-methyltransferase activity of TARBP1 in human cells, we first purified small RNA (<200nt) from TARBP1 KO and WT cells, and measured the Gm modification changes with high-performance liquid chromatography-mass spectrometry (HPLC-MS) (Extended Data Fig. 3b). As expected, the level of Gm modification in small RNA from TARBP1 KO cells showed a dramatic decrease, while no obvious changes were detected among other tRNA modifications, such as m^1^A (Extended Data Fig. 3c,d). Thus far, the Gm18 modification of tRNA in humans was identified in only two tRNA species (Gln and Ser) via high-throughput sequencing^14^. To further verify the tRNA substrates of TARBP1, we used biotin-labeled DNA probe to purify tRNA^Gln(TTG)^ and tRNA^Ser(TGA)^ from TARBP1 KO and WT cell lines and quantified Gm by HPLC-MS, respectively. As expected, the lack of Gm in tRNA^Gln(TTG)^ and a dramatic decrease of Gm in tRNA^Ser(TGA)^ were detected from TARBP1 KO cells (Fig. 3c,d,f). Notably, among Gm-containing tRNAs, 2’-O-methylation is not only present at position 18 but also at other positions, such as 34 and 39^20^. Thus, we purified tRNA^Phe(GAA)^ with a biotin-labeled DNA probe to quantify the level of Gm34 in the absence of TARBP1, and no significant change in Gm level was observed (Fig. 3e,f). Lastly, in order to understand the enzymatic activity and substrate specificity of TARBP1, we carried out *in vitro* assays by purifying recombinant wild-type (WT) and catalytic-dead mutant (Mut) of TARBP1 SpoU_methylase domain using prokaryotic expression system (Extended Data Fig. 3e,f). The recombinant WT or Mut protein was then incubated with substrate RNA probes including tRNA^Ser(TGA)^, tRNA^Gln^ ^(TTG/CTG)^, and tRNA^Phe(GAA)^, respectively. The catalytic activity of WT TARBP1 for the tRNA^Gln^ ^(TTG/CTG)^ oligos is approximately 2-fold higher than that for the tRNA^Ser^ ^(TGA)^ oligos (Fig. 3g,h). However, only a very low methylation activity was observed towards the tRNA^Phe(GAA)^ probes (Fig. 3g,h). Importantly, the enzymatic activity is completly inhibited when we substituted the nucleotides neighboring G18 (UGG to UGU, UGG to GGG) in the RNA substrate (Fig. 3i), suggesting TARBP1 catalytic activity is stringently dependent on UGG motif. Together, our findings suggest that TARBP1 is the Gm18 methyltransferase targeting tRNA^Gln^ ^(TTG/CTG)^ and tRNA^Ser^ ^(TGA/GCT)^ both in vitro and in vivo.

**Fig. 3:**
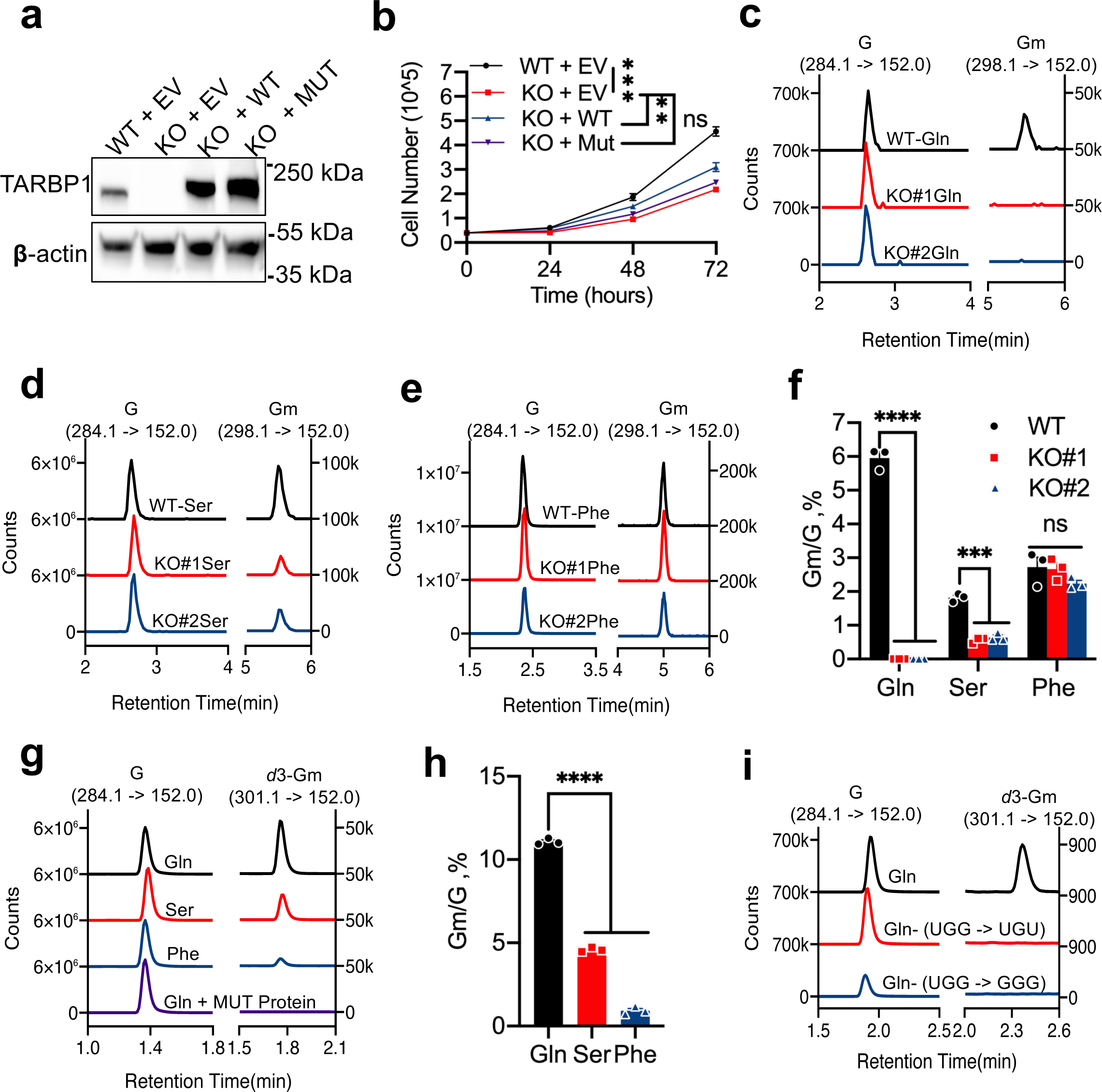
TARBP1 is the Gm18 methyltransferase targeting tRNA^Gln^ ^(TTG/CTG)^ and tRNA^ser (TGA/GCT)^ a, WT, TARBP1 KO, and KO cells rescued with WT or catalytically inactive mutant (Mut) TARBP1 protein were confirmed by western blot with TARBP1 antibody. β-actin was used as a loading control. b, Cell proliferation analysis of WT, TARBP1 KO, and WT or Mut rescue cells. c-f, LC-MS/MS analysis of Gm/G levels in purified tRNA^Gln(TTG)^ (c), tRNA^Ser(TGA)^ (d) and tRNA^Phe(GAA)^ (e) from the WT and TARBP1 KO cells. Quantitative analysis of LC-MS/MS analysis results (f). g, h, LC-MS/MS analysis of *d*3-Gm/G levels in substrate RNA probes (including tRNA^Ser(TGA)^, tRNA^Gln(TTG/CTG)^, tRNA^Phe(GAA)^) after being incubated with recombinant WT human TARBP1 protein in vitro (g). Quantitative analysis of LC-MS/MS analysis results (h). Mut TARBP1 protein group was used as a negative control quantification analysis. i, LC-MS/MS analysis of *d*3-Gm/G levels in tRNA^Gln^ probes after being incubated with recombinant WT human TARBP1 protein in vitro. We mutated the nucleotides neighboring G18 of tRNA tRNA^Gln^, including UGG mutation to UGU, and UGG mutation to GGG. **P<0.01, ***P<0.001, ****P<0.0001, ns: not significant. Student’s t-test. b, Data represented as mean ± SD (n=3). f, h, Data were presented as mean ± SEM (n=3).

### TARBP1 depletion decreases the stability of tRNA^Gln^ ^(TTG/CTG)^ and tRNA^Ser(TGA/GCT)^

To unveil the molecular mechanism underlying the tumor-suppressive role observed in TARBP1-KO cells, we employed several different methods to explore the Gm18 functions of modified tRNAs. Firstly, we analyzed the levels of tRNA^Ser(TGA)^ and tRNA^Gln(CTG)^ by performing a Northern blot assay and observed a notable reduction in those Gm18-modified tRNAs in TARBP1 depleted cells compared with WT cells (Fig. 4a). Secondly, we further validated the stability of those tRNAs by tRNA-seq and identified a remarkable increasing levels of tRNA-derived RNA fragments (tRFs) derived from tRNA^Ser(TGA/GCT)^ and tRNA^Gln(TTG/CTG)^ in TARBP1 KO RNA samples (Fig. 4b). As shown in Extended Data Fig. 4a, there are five structural types of tRFs, including 5’-tRNA halves, 3’-tRNA halves, 5’-tRFs, 3’-tRFs, and i-tRFs, and one of the structural categories of i-tRFs begin at position 18. Interestingly, further analysis of tRNA-seq confirmed that those increased tRFs in the TARBP1 KO group mainly belong to i-tRFs surrounding position 18 (Fig. 4c and Extended Data Fig. 4b). Taken together, the above data demonstrated that TARBP1 deficiency caused loss of the Gm18 tRNA modification in tRNA^Gln^ ^(TTG/CTG)^ and tRNA^Ser^ ^(TGA/GCT)^, thus reducing the stability of these tRNAs, in particular tRNA^Gln^ ^(CTG)^.

**Fig. 4:**
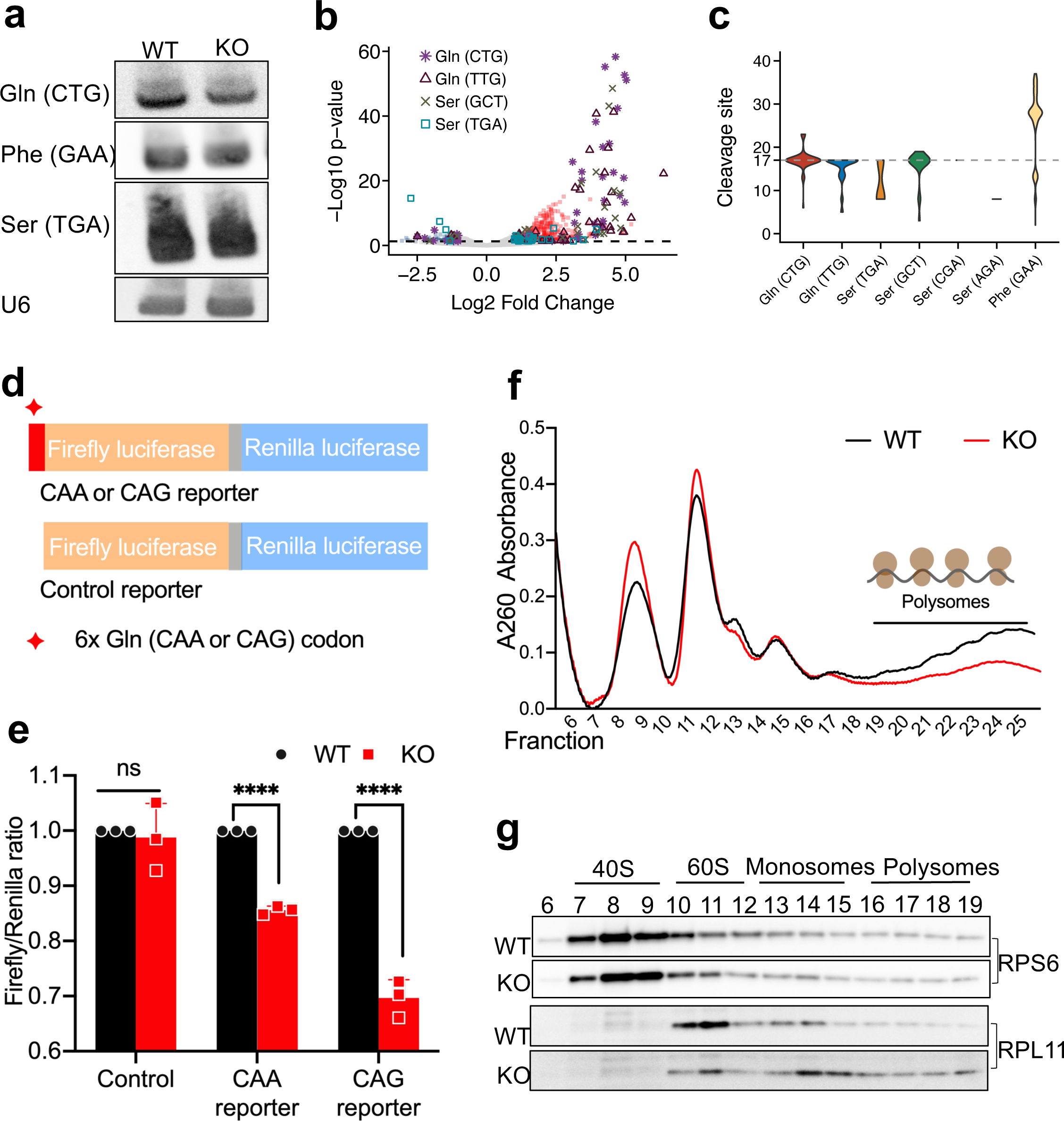
TARBP1 affects global translation possibly by regulating the structural stability of Gm18-modified tRNAs. a, Gm18-modified tRNA levels in WT and TARBP1 KO groups were measured via northern blot. b, tRNA fragments (tRFs) in WT and TARBP1 KO RNA samples were analyzed by tRNAseq. c, Analysis of tRF initiation sites of the increased tRFs upon loss of TARBP1. d, Scheme of reporter vector enriched with 6 x CAA or 6 x CAG codons. tRNA-Gln (TTG) reporter plasmid was gained by inserting CAACAACAACAACAACAA after the start codon of F-Luc, tRNA-Gln (CTG) reporter plasmid was gained by inserting CAGCAGCAGCAGCAGCAG after the start codon of F-Luc. e, Translation efficiency of firefly luciferase fused with 6 x CAA or 6 x CAG codons in WT and TARBP-KO cells was revealed by a reporter assay. Firefly light units were normalized to Renilla luciferase, and the empty vector was set to 1. ****P<0.0001, ns: not significant. Student’s t-test. Data represented as mean ± SD (n=3). f, g, Global translation in WT and TARBP1 KO group were detected by polysome profiling (f), and western blot analysis with RPS6 and RPL11 antibodies confirmed ribosome fractionation (g).

### TARBP1 enhances glutamine metabolism via translation regulation

The tRNA is essential for protein synthesis, which can recognize mRNA codons and bridges corresponding amino acids. Growing evidences show that tRNA modifications are crucial for translation regulation^20–25^. Hence, we wondered whether Gm18 modification regulates mRNA translation via modified tRNAs. The dual luciferase reporter plasmid in which we inserted 6 x Gln (CAA) or 6 x Gln (CAG) codons after the start codon of firefly luciferase was used to measure protein translation in WT and KO cells. We observed that the expression of firefly luciferase in WT cells was higher than KO cells, indicating that Gm18 modification may impact the expression of the proteins frequently using modified tRNAs (Fig. 4d,e). Polysome profiling was also performed to measure global protein synthesis and it was revealed a striking reduction of polysome fractions in TARBP1-depleted cells compared with WT cells, suggesting that TARBP1 is critical for global translation in cancer cells (Fig. 4f,g). We next performed ribosome profiling sequencing (Ribo-seq) to identify the mRNA translation changes in cells depleting TARBP1 and control cells (Extended Data Fig. 5a-g). We identified 2259 differentially translated genes with high confidence (Ribo transcripts per million > 1, RNA transcripts per million > 5, fold change >= 1.5, and p-value < 0.05). Among them, there are 1167 genes with decreased translation efficiency and 1092 genes with increased translation efficiency in the TARBP1 KO group compared with the WT group (Fig. 5a). We next asked whether the tRNA Gm18 modification mediated by TARBP1 can regulate mRNAs translation through Gm18 tRNA-decoded codon usage. It was reported that decreased tRNA abundance is expected to increase codon-specific ribosome dwell time^26^. Further analysis of Ribo-seq uncovered that the higher CAG codon occupancy was observed in KO sample compared with WT control, and the translation efficiency of mRNAs with higher CAG codon frequency were more inclined to be down-regulated upon TARBP1 deletion (Extended Data Fig. 4c,d).

**Fig. 5:**
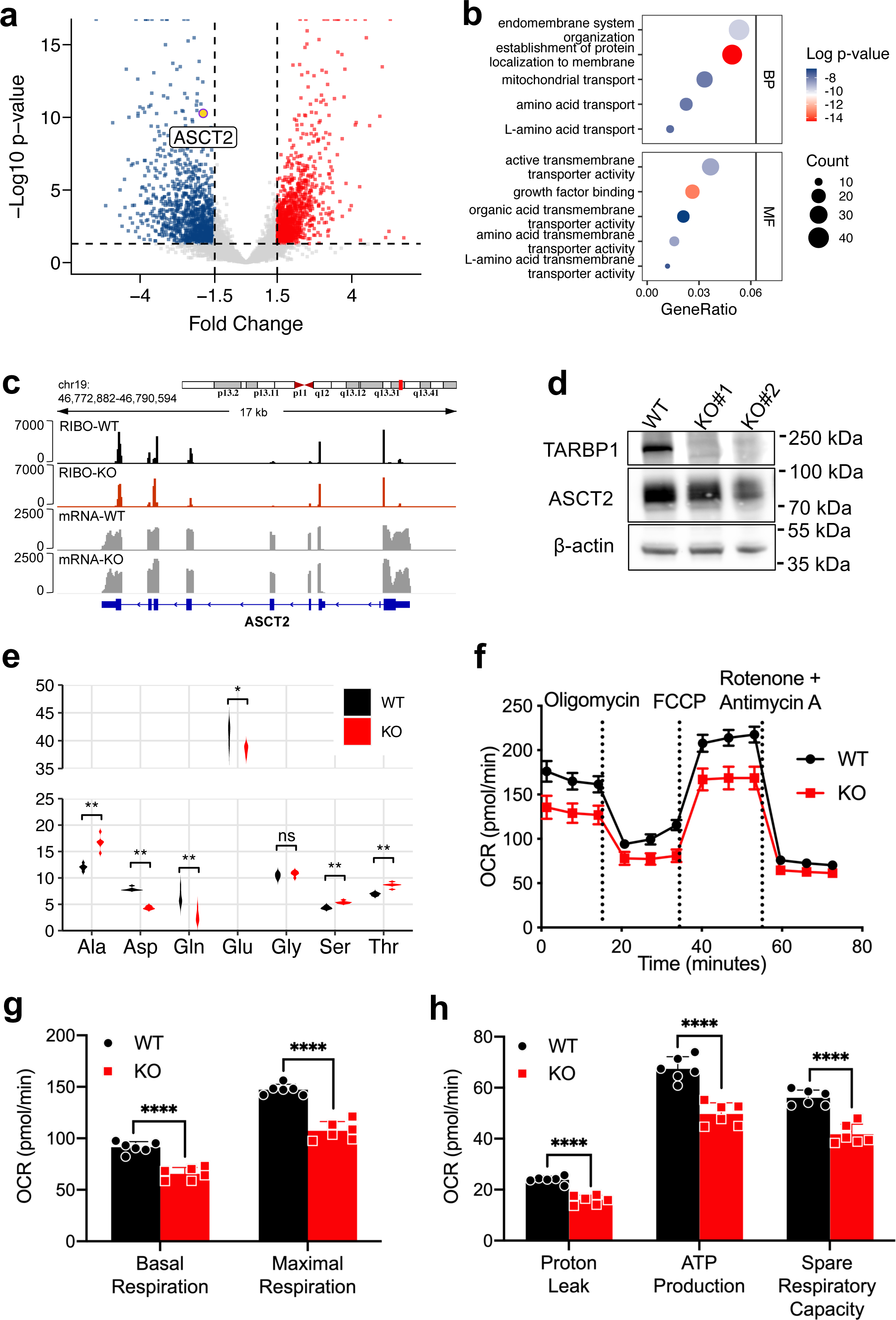
TARBP1 enhances glutamine metabolism via translation regulation. a, Volcano plot depicting down-regulated (blue) and up-regulated (red) genes in TARBP1 KO cells compared with WT cells in Ribo-seq. b, Gene Ontology analysis of reactive pathway enrichment using the TE downregulated genes upon TARBP1 deficiency. c, Track plot showing translational changes of ASCT2 in TARBP1 KO cells compared with control cells. d, Western blot analysis confirming ASCT2 expression in control and TARBP1 KO cells. β-actin was used as a loading control. e, LC-MS/MS analysis of intracellular amino acid concentration in TARBP1 KO cells compared with WT cells, and six independent samples were used to plot. f-h, Oxygen consumption rate of WT and TARBP1 KO cells measured by using a Seahorse XF96 analyzer (f). Quantification of the basal and maximal OCR for the indicated cells (g). Quantification of the proton leak, ATP production, and spare respiratory capacity OCR for the indicated cells (h). ****P<0.0001. Student’s t-test. Data represented as mean ± SD (n=6).

Notably, Gene Ontogeny analysis revealed that mRNAs with down-regulated translation efficiency in the TARBP1 KO cells are enriched in the pathway of amino acid transport (Fig. 5b). Moreover, the translation efficiency of ASCT2 (also known as SCL1A5) mRNA was severely suppressed in KO cells (Fig. 5c). Consistently, western blot further confirmed that the protein level of ASCT2 was significantly decreased in TARBP1-depleted cells (Fig. 5d). Given that ASCT2 was demonstrated as a major transporter of glutamine^27, 28^, we speculated that the oncogenic effects of TARBP1 is partially due to enhanced tumor glutamine metabolism, which is in line with our screening results (Fig 1). Consistent with this hypothesis, lower levels of glutamine and glutamate were detected in TARBP1 KO cells compared with WT cells (Fig. 5e). Additionally, metabolic flux analysis uncovered that TARBP1 deficiency significantly suppresses oxidative phosphorylation in cancer cells (Fig. 5f-h). Taken together, these data provide further evidences for the biological function of TARBP1-mediated Gm18 tRNA modification in glutamine metabolic reprogramming.

### TARBP1 and ASCT2 are co-expressed in liver cancer and associated with HCC patient prognosis

To further verify the results observed in cell model, we determined to investigate the clinical significance of TARBP1 in HCC clinical samples. Consistent with our cell model results, the IHC staining results showed that both TARBP1 and ASCT2 were remarkablely upregulated in tumors compared with paratumors (Fig. 6a,b). More importantly, the expression level of ASCT2 in HCC tissues was positively correlated with the expression of TARBP1: samples with higher TARBP1 expression exhibited high ASCT2 levels, while samples with lower TARBP1 expression exhibited low ASCT2 levels (Fig. 6c and Extended Data Fig. 6a). Additionally, the patients with higher expression levels of TARBP1 or ASCT2 have worse overall survival (Fig. 6d) and higher cumulative recurrence rate after surgery (Fig. 6e). Interestingly, compared with other tRNA species, higher tRNA(Gln) levels were observed in hepatobiliary cancers (Extended Data Fig. 6b). In summary, our study prove that TARBP1 is a novel oncogene through mediating tRNA modification and stabilizing a small subset of tRNAs (in particular Gln-tRNA), which sustains the efficient translation of ASCT2, in turn enhances the uptake of glutamine and promotes the tumorigenesis (Fig. 6f)

**Fig. 6:**
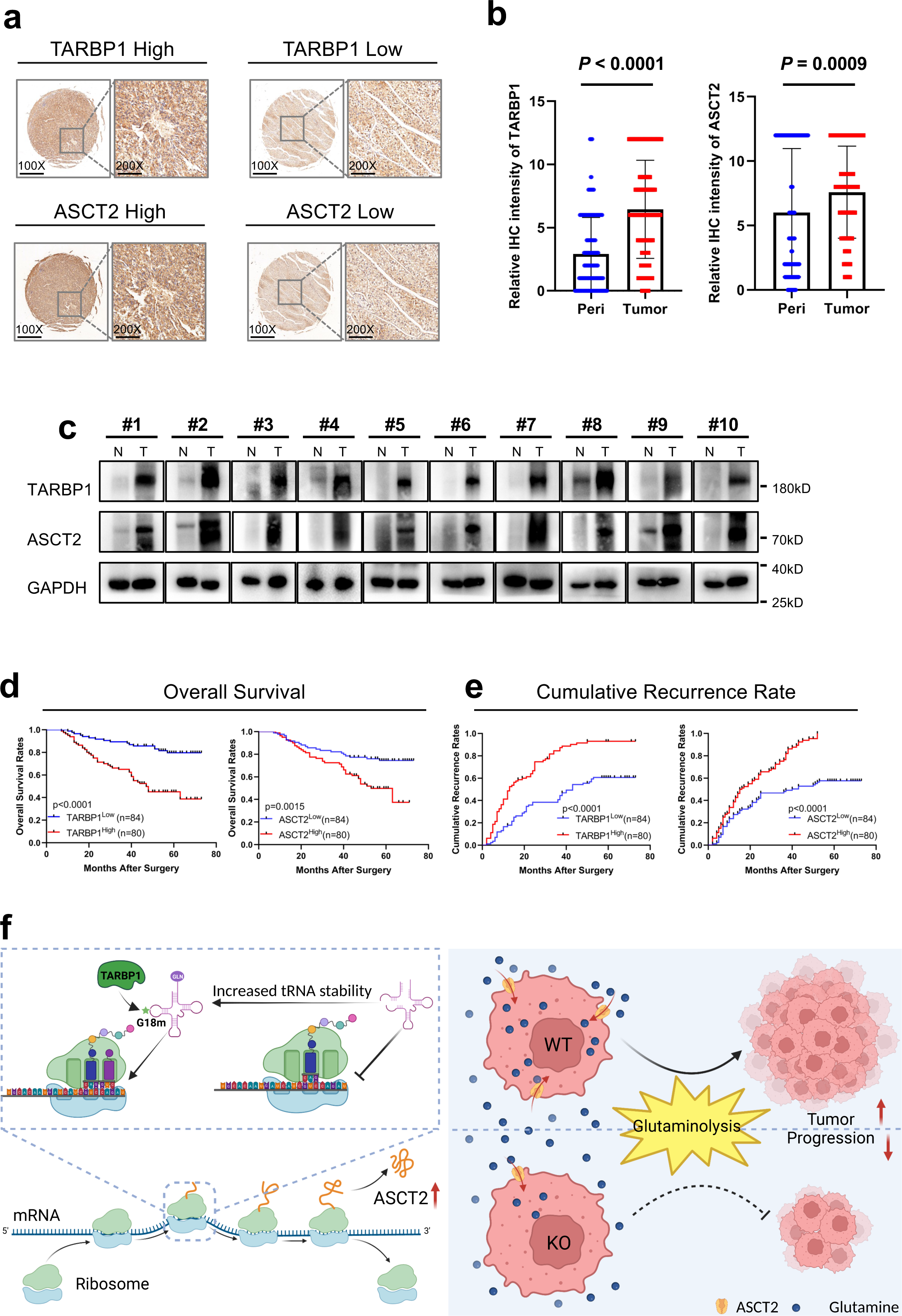
The levels of TARBP1 and ASCT2 correlate and are associated with HCC patient prognosis. a, Representative IHC staining images showing TARBP1(top) and ASCT2 (Bottom) in HCC tissues, respectively. b, Quantitative IHC staining intensity of TARBP1(left) and ASCT2 (right) in HCC tissues and para tissues. Peri: para tissues. Student’s t-test. Data represented as mean ± SD (n= 164). c, Western blot analysis showing TARBP1 and ASCT2 expression in HCC tissues and normal tissues, respectively. T: HCC tissues, N: normal tissues. GAPDH was used as a loading control. d, Analysis of overall survival in patients with low and high expression of TARBP1(left) and ASCT2 (right). e, Analysis of cumulative recurrence rate in patients with low and high expression of TARBP1(left) and ASCT2 (right). f, Summary model for the critical roles of TARBP1 in HCC progression through glutamine metabolic reprogramming. TARBP1 is a novel oncogene through mediating tRNA modification and stabilizing a small subset of tRNAs (in particular Gln-tRNA), which sustains the efficient translation of ASCT2, in turn enhances the uptake of glutamine and promotes the tumorigenesis

## DISCUSSION

In current study, we demonstrated that TARBP1 is novel glutamine metabolism regulator through CRISPRi-Cas9 screening. Importantly, up-regulation of TARBP1 - commonly via gene amplification-is frequently detected in multiple types of human cancers and is disfavorable for patient survival. In line with the observation, knockout of TARBP1 significantly suppresses cancer cell proliferation, arrests cell cycle progression, inhibits colony formation and blocks in vivo tumor formation, which highlights TARBP1 as a possible oncogene in many cancers. Furthermore, our in vitro and in vivo assay showed that TARBP1 mediates Gm18 methylation targeting tRNAGln (TTG/CTG) and tRNASer (TGA/GCT), and Gm18 modification can enhance the stability of modified tRNAs. TARBP1 depletion leads to a substantial decrease in the levels of tRNAs that harbor the Gm18 modification, and globally decreases mRNA translation. Interestingly, Ribo-Seq showed that a subset of genes enriched in the pathway of transporter are specifically affected upon TARBP1 knockout. Of note, the essential glutamine transporter ASCT2 is the major downstream target of TARBP1. Overall, these results reveal the underlying molecular and cellular mechanism of TARBP1 oncogenicity that involves increased mRNA translation of ASCT2, which fuels the tumorigenesis via enhancing glutamine metabolism.

### The catalytic specificity of TARBP1 SpoU domain

TARBP1 was initially identified as one of the cellular factors that bind with both high affinity and marked specificity to TAR RNA loop sequences and are involved in the regulation of HIV-1 transcription^29–32^. Of note, TAR RNA region highly resembles the D-loop patch of tRNA-Gln/Ser, with the respects of both sequence and secondary structure^33^. A very recent study showed that TARBP2 recruits the RNA methyltransferase FTSJ3 to decorate multiple HIV RNA sites with 2-O-methylation which enables the HIV RNA to escape from immune recognition, though it’s still not clear about whether TARBP1 is also hijacked by HIV virus in this context^34^. Interestingly, our *in vivo* and *in vitro* assays demonstrated that TARBP1 SPOUT domain shows quite stringent substrate preference and only catalyzes a small subset of tRNAs, including tRNA-Gln/Ser, which have specific signatures in sequence context and/or structures. Consistent with this hypothesis, we found that the methylation activity is largely abolished upon the mutagenesis/substitution of neighboring G into A/C in synthesized substrates. Ultimately, a co-crystal structure of TARBP1 (ideally the full-length protein) with its RNA substrates will be helpful for unconvering the detailed mechanism of TARBP1’s specificity towards tRNA-Gln/Ser.

### Regulatory role of tRNA Gm18 modifications

Methylation of riboses at 2-OH group is one of the most common and widespread RNA modifications found in many RNA species, including ribosomal RNA, transfer RNA, miRNA and messenger RNA^35^. In particular, tRNA molecules from all three domains of life are 2-O-methylated in multiple residues and the methylation patterns are commonly conserved while also evolving from bacteria to mammals^14^. Both in bacteria and yeast, Gm18 modification is deposited in a large category of tRNAs, while depletion of Gm18 methyltransferase doesn’t have significant effect on cell growth or on translation efficiency under normal condition^14, 17^. Moreover, previous studies demonstrated that Gm18 is a global immune inhibitory tRNA modification and contributes to tRNA recognition by innate immune cells^36–38^. These observations raised the hypothesis that this modification may mainly function as an immune escape mechanism but not in mRNA translation. Unexpectedly, our results uncovered that the loss of Gm18 in tRNA causes a catastrophic effect towards tRNA stability and cell growth in cancer cells of mammals, though only limited types of tRNA (tRNA-Gln/Ser) still bear this modification. Considering that Gm18 interacts with U54 of T-loop in structured tRNA, it may play critical role for maintaining the tRNA tertiary structure and tRNA stability^39^. Indeed, it was reported that the complete lack of Gm18 in rat hepatoma cell tRNASer (codon IGA) causes noticeable changes in elution profiles of this RNA compared with Gm18-modified form, indicating that the tRNA tertiary structure is affected upon loss of this modification. Besides the potential involvement in tRNA structure maintaining, it’s also possible that G18m stabilizes the tRNA by preventing tRNA from attacking and degradation by endonuclease. Indeed, a myriad of recent studies have proved that the 2-O-methylation is frequently employed by RNA virus to evade from degradation system of host during infection^40, 41^.

### tRNA modifications and cancer

The tRNA is decorated by various types of chemical modifications during its biogenesis. Interestingly, the modifications can occur in the functionally important sites of all tRNAs (Extended Data Fig.3a) and have enormous implications in modulating gene expression, particularly gene-specific translation^20–22^. For example, modifications in wobble positions can enhance base pairing and increase the translation speed and fidelity of codon-specific genes^23^. Modifications outside the anticodon loop also regulate gene expression by modulating tRNA biogenesis, including tRNA processing, cleavage, and structural stability^24, 25^. Our discovery here unexpectedly uncovered the important roles of TARBP1 and G18m modification in tumorigenesis through regulating glutamine metabolism. Considering the close connection between availability of amino acids and translation, it will be appealing to speculate that there will be more similar mechanism awaiting further exploration.

In summary, our study prove that TARBP1 is a novel oncogene through mediating tRNA modification and stabilizing a small subset of tRNAs (in particular Gln-tRNA), which sustains the efficient translation of ASCT2, in turn enhances the uptake of glutamine and promotes the tumorigenesis. These findings pave the new avenues of developing therapeutic strategies against TARBP1 for effective cancer treatment.

## MATERIALS AND METHODS

### Plasmid construction

To re-express TARBP1 protein into KO cells, TARBP1 cDNA was cloned into the pCDH-EF1-Luc2-P2A-copGFP retroviral vector. Then the QuikChange Site-Directed Mutagenesis Kit (Stratagene, 200518) was used to generate TARBP1 catalytically inactive mutant (Mut, amino acids position at 1482, Arg to Ala). Truncated WT or Mut ORF DNA of TARBP1 (amino acids position 1425 ∼ 1621) was inserted into the pET32a with 6xHis tag expression vector to express truncated TARBP1 protein. Two TARBP1-specific 20nt small guide RNA (sgRNA) sequences designed to target the first exon of TARBP1 were cloned into pL-CRISPR.EFS.GFP plasmid separately.

sgRNA#1: TGGCGCGGGCGCTCGGCAAA

sgRNA#2: GCCGCTACTGGAGCGCGTGG

### Cell culture

Cells were cultured in Dulbecco’s Modified Eagle Medium (DMEM, Gibco) containing 10% fetal bovine serum (FBS, BI) and 1% penicillin-streptomycin (Thermo Fisher Scientific) unless otherwise specified, and were maintained in a 37°C humidified incubator with 5% CO2.

### Cell proliferation, colony formation, and migration

For cell proliferation, 3 × 10^5^ cells were seeded in one well of a 12-well plate on day 0 and counted on day one until day 3 with cell counters (Thermo Fisher Scientific). For colony formation, 1000 cells were seeded in a 6-cm dish and cultured for nine days. Cells were then fixed with 4% formaldehyde (Beyotime) and stained with 0.1% crystal violet solution (Solarbio). Next, the number and size of Cell colonies were measured. For the wound-healing assay, 6 × 10^5^ cells were seeded in one well of a 6-well plate, and cells were cultured in DMEM with 10% FBS until cells at approximately 90-100% confluency and were then cultured in DMEM with 2% FBS overnight. Scratch wounds were created by a pipette tip and captured with a microscope at 24 hours later.

### Apoptosis and cell cycle analysis

For apoptosis analysis, a 6-well plate of cells at approximately 80-90% confluency was harvested by treatment with trypsin without EDTA (Gibco). The staining was performed by the use of an Annexin V-FITC/PI Apoptosis Detection Kit (Vazyme), and the apoptotic cells were detected by flow cytometry analysis (BD FACSCanto). For cell cycle analysis, a 6-well plate of cells at approximately 80-90% confluency was harvested by treatment with trypsin without EDTA and were then fixed in cold 70% ethanol for 2 hours at 4°C. The staining procedures were performed by the use of a Cell Cycle and Apoptosis Analysis Kit (Yeasen) and the cell cycle were detected by flow cytometry analysis (BD FACSCanto).

### Recombinant protein purification

To express recombinant wild-type and Mut TARBP1 proteins, pET32a TARBP1-WT or TARBP1-Mut vector was transformed into BL21 bacteria, respectively. The bacterial cells were cultured in LB medium at 37°C until OD600 reached 0.6–1. Protein expression was induced by the use of IPTG with 0.1 mM final concentration at 16°C for 16 hours. The bacteria pellets were collected by centrifugation at 2000rpm at 4°C for 10 min and then lysed with the lysis buffer (300 mM NaCl, 25 mM Tris pH 7.5, 10% Glycerol, and 0.5% NP-40). Clarify cell lysates by centrifuging at 20,000rpm at 4°C for 40 min and purify recombinant protein via HisPur Ni-NTA Agarose.

### Generation of TARBP1 KO cell lines and Rescue experiments

To generate TABRP1 knockout cells, 2 μg of sgRNA plasmid was transfected to PLC/PRF/5 cells using Lipofectamine 3000 (Invitrogen). After 2 days’ culture in DMEM medium, cells were screened according to fluorescence and isolated into monoclons by flow cytometry. After 3 weeks’ culture, colonies were screened out and confirmed by western blot with TARBP1 antibody.

To generate viruses, the empty vector, flag-TARBP1 WT, or flag-TARBP Mut plasmid was co-transfected with pMD2.G and PsPAX2 plasmids into 293T cells, respectively. PLC/PRF/5 WT cells were infected with virus generated by empty vector, whereas KO cells were infected with virus generated by empty vector, flag-TARBP1 WT, or flag-TARBP Mut plasmid separately. After 7 days’ selection with 2 μg/ml puromycin, cells were confirmed by western blot with TARBP1 antibody.

### CRISPRi/Cas9 library screening

For generating stably expressed Cas9 in B16 melanoma cells, the vector containing the coding sequence of both Cas9 and fluorescence protein was introduced into cells via lentiviral transduction. The transduced cells were then screened according to fluorescence and sorted by flow cytometry. For individual gene knockdown in B16 cells, sgRNAs were stably transfected into B16 cells with Cas9 stably expression. Subsequently, Cells were selected with 2 μg/mL of puromycin for 7 days for screens. Infected cells were cultured in glutamine - basic medium (2mM glutamine, 10% FBS, 1% penicillin-streptomycin) or complete medium (10mM Glucose, 1mM pyruvate, 2mM glutamine, 10% FBS, 1% penicillin-streptomycin) for 48 hours. Dead cells cultured in glutamine - basic medium and cells cultured in complete medium were harvested to extract genomic DNA (gDNA) and amplify sgRNA from 5 μg gDNA using 2 X PCR Master Mix. The prepared libraries were then subjected to massive parallel high-throughput sequencing by Berry Genomics. The CRISPRi/Cas9 screening data analysis was performed by MAGeCK(v0.5.9.5) as previously described^18^. MAGeCK RRA method was used to evaluate the abundance of sgRNAs and identify gene hits.

### Endogenous tRNAs purification by biotin-labeled DNA probes

Using streptavidin T1 beads (Invitrogen), we purified the endogenous tRNA^Ser(TGA)^, tRNA^Gln^ ^(TTG)^, and tRNA^Phe(GAA)^ with biotin-labeled DNA probes. First, 200 pmol of total RNAs were mixed with 2,000 pmol of specific biotinylated DNA probes in 0.3 volumes of hybridization buffer (250 mM HEPES pH 7, 500 mM KCl) and the mixtures were incubated at 90°C for 7min, then cooled down slowly to 25°C over 3.5 hours. Subsequently, the hybridization mixtures were mixed with beads in IP buffer (150 mM NaCl, 50 mM Tris, pH 7.9, 0.1% NP40) and incubated at 4°C for 1 hour. Finally, the oligonucleotide-conjugated beads were washed with IP buffer for five times and heated at 70°C for 3min to elute tRNA for QQQ analysis.

For tRNA^ser(TGA)^: 5’ Biotin-CGCCTTAACCACTCGGCCACGACTAC

For tRNA^Gln^ ^(TTG)^: 5’ Biotin-AGTGCTAACCATTACACCATGGGACC

For tRNA^Phe(GAA)^: 5’ Biotin-CGCTCTCCCAACTGAGCTATTTCGGC

### Assays for 2’-O methyltransferase activity in vitro

In vitro methyltransferase assay was performed at 37°C for 2 hours in 30 ul reaction mixtures containing 1 μg RNA probes, 2 μg fresh recombinant protein, 0.1 mM *d*3-SAM, 50 mM Tris–HCl pH 7.5, 10% glycerol. The resultant RNA was then digested with nuclease P1 (NEB) and antarctic phosphatase (NEB) and subjected to LC-MS/MS analysis to quantify different nucleosides.

tRNA^Ser^ ^(TGA)^: GCCGAGUGGUUAAGGC

tRNA^Gln^ ^(TTG/CTG)^: GUGUAAUGGUUAGCAC

tRNA^Phe^ ^(GAA)^: AGACUGAAGAUCU

tRNA^Gln^ ^(Mut^^1)^: GUGUAAUUGUUAGCAC

tRNA^Gln^ ^(Mut^^2)^: GUGUAAUGGGUAGCAC

### Polysome profiling

Two 15-cm plates of PLC/PRF/5 cells prepared for each sample (WT and TARBP1 KO) were treated with 100 μg/ml Cycloheximide (MedChemExpress) and incubated for 8 min at 37 °C. Cells were lysed on ice for 10min in 400uL lysis buffer (20 mM Tris– HCl pH 7.5, 150 mM NaCl, 5 mM MgCl_2_, 1 mM DTT, 100 μg/ml Cycloheximide, 1% Triton X-100). The supernatant was collected via centrifugation at 12 000 g for 10 min at 4 °C, and its’ absorbance at 260nm was measured. The ribosomes with equal A260 in each group were loaded onto a sucrose gradient (10–50%) buffer prepared using a gradient station and centrifuged in an SW41Ti rotor (Beckman) at 4°C for 4 hours at 28,000 rpm. Polysome profiles were then generated and analyzed by Gradient Station.

### tRNA Seq

Total RNA samples isolated from PLC/PRF/5 cells with or without TARBP1 depletion were first subjected to small RNA purification using the Zymo Research RNA Clean & Concentrator-25 kit (Zymo). The isolated small RNAs were heated at 75°C for 2 min to denature tRNA and then incubated with ALKB-D135S protein in buffer containing 300 mM KCl, 2 mM MgCl_2_, 50 µM of (NH4)2Fe(SO4)2·6H2O, 300 µM 2-ketoglutarate (2-KG), 2 mM l-ascorbic acid, 50 µg/ml BSA, 50 mM MES buffer (pH 5.0) for 2 hours at room temperature. The demethylated RNAs were purified by the Zymo Research RNA Clean & Concentrator-25 kit and then treated with PNK. RNA library was prepared by VAHTSTM Small RNA Library Prep Kit for Illumina® (Vazyme).

### Ribo-seq

Two 15-cm plates of PLC/PRF/5 cells with approximately 80-90% confluency were used per group. Cells were incubated with 100 μg/ml Cycloheximide for 8 min at 37°C and were then lysed on ice for 10min in 400uL lysis buffer (20 mM Tris–HCl pH 7.5, 150 mM NaCl, 5 mM MgCl_2_, 1 mM DTT, 100 μg/ml Cycloheximide, 1% Triton X-100). The supernatant was collected via centrifugation at 12 000 g for 10 min at 4 °C, and its’ absorbance at 260nm was measured. The ribosomes with equal A260 in each group were reserved for RNA sequencing. Added 15U RNase1 for each 1OD of lysate, and incubated the reactions at 22°C for 40 min with gentle mixing. Ribosome-protected fragments (RPFs) were collected by MicroSpin S-400 columns (Cytiva) and purified by Zymo Research RNA Clean & Concentrator-25 kit. The RPFs were then separated with 15% polyacrylamide TBE–urea gel, excised the gel slices between 17 and 34 nucleotides and recovered RNA from PAGE gel by ZR small-RNA™ PAGE Recovery Kit (Zymo). Purified RNA was treated with PNK, and the RNA library was prepared by VAHTSTM Small RNA Library Prep Kit for Illumina®.

### Luciferase Reporter Assay

PmirGlo vector containing the luciferase of firefly (F-Luc) and Renilla (R-luc) was used to perform Dual luciferase assays. The activity of F-Luc normalized by R-Luc activity was to evaluate the reporter translation efficiency. CAA or CAG reporter plasmid was gained by inserting CAACAACAACAACAACAA or CAGCAGCAGCAGCAGCAG after the start codon of F-Luc. 1μg of empty vector, CAA reporter plasmid, or CAG reporter plasmid were transfected into TARBP1 KO and WT PLC/PRF/5 cells, respectively. After 48 hours of transfection, the luciferase expression was detected by a Dual-Glo Luciferase Assay System (Promega).

### Northern blot

For northern blotting, 10 μg of total RNA was mixed with 3 μL of 2xGel Loading Buffer and heated at 70°C for 5 min. After pre-run of the 15% TEB-Urea gel, samples were subjected to the gel for electrophoresis separation. Next, RNA was transferred onto a membrane and subsequently crosslinked with UV. The membrane was further incubated with the corresponding RNA probe and rotated at 42°C overnight. Membranes were then incubated with Streptavidin-HRP (Invitrogen) for 45 min at room temperature and were finally subjected to a phosphoimager following several times washes.

U6 snoRNA: GCAGGGGCCATGCTAATCTTCTCTGTATCG

tRNA^Gln(CTG)^: GTTCCACCGAGATTTGAACTCGG

tRNA^Ser(TGA)^: CGCCTTAACCACTCGGCCACGACTAC

tRNA^Phe(GAA)^: CGCTCTCCCAACTGAGCTATTTCGGC

### Western blot

For western blotting, a six-well plate of PLC/PRF/5 cells at approximately 80-90% confluency was lysed with 100 μl RIPA buffer on ice. Cell lysates were separated by 4-20% SDS-PAGE, and the protein was then transferred to 0.4 μm polyvinylidene fluoride (PVDF, Merck Millipore) membrane. After blocking with 5% milk in TBST, the membranes were then incubated with the corresponding antibodies overnight at 4°C. Membranes were incubated with secondary HRP-conjugated antibody (proteintech) at room temperature for 1 hour following three times washes with TBST and were washed three times again before detected by the phosphoimager.

The corresponding antibodies included Monoclonal rabbit anti-TARBP1 antibody with 1: 1000 dilution (abcam), Monoclonal rabbit anti-ASCT2 antibody with 1: 3000 dilution (proteintech), Polyclonal rabbit anti-RPS6 antibody with 1: 1000 dilution (proteintech), Polyclonal rabbit anti-RPL11 antibody with 1: 1000 dilution (proteintech), Monoclonal rabbit anti-GAPDH antibody with 1: 5000 dilution (proteintech) and Monoclonal rabbit anti-β-actin antibody with 1: 3000 dilution (SAB Signalway Antibody).

### Seahorse

The oxygen consumption rate (OCR) of WT and TARBP1 KO cells was measured by using the XF96 extracellular flux analyzer (Agilent). 30000 cells were seeded onto one well of mini plates, and then maintained in a 37°C incubator with 5% CO2 overnight. Subsequently, cells were washed three times with assay medium and maintained in an incubator without 5% CO2 for 1 h. The OCR of each wall was measured when oligomycin (1 μM), carbonyl cyanide 4-(trifluoromethoxy) phenylhydrazone (FCCP; 1.5 μM), and rotenone/antimycin A (0.5 μm) were sequentially injected.

### Animal studies

A xenograft experiment was performed to evaluate the tumor-suppressive role of TARBP1. 1x10^6^ cells mixed with equipped volume of Matrigel (Corning) were injected into the right flank of nude mice. Subcutaneous tumor formation was measured on day 7 and then every other day until the maximal tumor size reached 2000 mm^3^. The tumor volume is calculated as the product of length x width^2^. At end-point, the tumors were collected to measure tumor weight. All mice were bred under specific-pathogen-free (SPF) conditions at the Animal Experiment Center of SUSTech University.

### Clinical specimens

Tumor specimens and para-tumor tissues used in this study were randomly collected from 164 consecutive patients with HCC who underwent curative resection at the Liver Cancer Institute of Fudan University (Shanghai, China). The surgical time was from July 2015 to December 2017 and the follow-up information was available from November 2015 to August 2021. The survival time was 7∼73 months with a median survival time of 23 months. Ethical approval was obtained from the Zhongshan Hospital Research Ethics Committee, and written informed consent was obtained from each patient.

HCC samples were reviewed histologically by hematoxylin and eosin staining, and representative areas of tumor and para-tumor were pre-marked in the paraffin blocks by professional pathologists. Duplicates of 1-mm-diameter cylinders from HCC samples were embedded in a new wax block. Tissue microarray blocks containing 164 cylinders were constructed, and sections 4 μm thick were placed on slides coated with 3-aminopropyltriethoxysilane.

Immunohistochemistry of the tissue microarray was performed using Dako REALTM EnVisionTM Detection System, according to the manufactory’s instructions. Briefly, microarray was dewaxed and then hydrated. After microwave antigen retrieval, the chips were incubated in blocking solution. Then the TMA were stained with the ASCT2 or TARBP1 antibody and a further incubation with second antibody reagent. The immunostaining was developed using the substrate 3,3’-diaminobenzidine (DAB) and the nuclei were counterstained with hematoxylin. After staining, the microarray was scanned using Aperio XT Slide Scanner. And the staining results were scored by two professional pathologists.

### Pre-processing of sequencing data

Ribo-seq, tRNA-seq, and RNA-seq data were pre-processed as described previously^42^. In brief, adapters and low-quality bases were trimmed with TrimGalore (version 0.6.7). Clean data were first mapped to rRNA sequences from SILVA rRNA database (release 138)^43^ and repetitive sequences from RepBase (v27.04)^44^ by Bowtie (version 1.3.1)^45^ with -v 1 --best, and the remaining unaligned reads were demultiplexed for further analysis.

### Ribo-seq and RNA-seq data analysis

Processed reads were aligned to the reference genome (GRCh38) by the STAR (version 2.7.10a)^46^, allowing for at most two mismatches. StringTie (v2.1.7)^47^ was used to calculate the raw read count. Both read counts of Ribo-seq and RNA-seq were converted to TPM (transcripts per million). Quality control was performed by Ribotoolkit^48^. Genes with sufficient expression level (Ribo TPM > 1 and RNA TPM > 5) were subjected to further analysis. Translation efficiency (TE) and the differentially-TE genes (DTEGs) were calculated by RiboDiff (v0.2.2)^49^. GO term analyses were performed by Metascape (https://metascape.org/gp/index.html). GraphPad Prism software and R was used for data presentation.

### tRF analysis

Trimmed reads in FASTQ format were given as input to MINTmap^50^ for identifying and quantifying reads associated with tRFs. The sample normalization and differential expression was performed using the DESeq2 (version 1.32.0)^51^.

## ACKNOWLEDGMENTS

We would like to thank all members in Hao Chen’s Lab for their help and advice in experimental design. We thank Prof. Ruilin Tian for sharing the CRISPRi/Cas9 library screening vectors and all kind suggestions. The authors would also like to acknowledge the technical support from Hua Li and Lin Lin at SUSTech CRFT. This work was supported by Center for Computational Science and Engineering at Southern University of Science and Technology.

## AUTHOR CONTRIBUTIONS

H.H.M and H.C. designed and conceived the experiments. J.B.C., J.W., and L.Y.W. provided clinical samples and performed pathological analysis. Y.Y.Z. and X.Y.S. performed experiments. R.Q.W. assisted in CRISPRi/Cas9 library screening. Y.G. helped prepare figures. M.G.X. and N.J.O assisted in polysome profiling. L.Q., Y.C.W. and H.C. analyzed the data. X.Y.S. and H.C. wrote the manuscript. All authors have read and approved the final manuscript.

---

**Extended Data Fig. 1:**
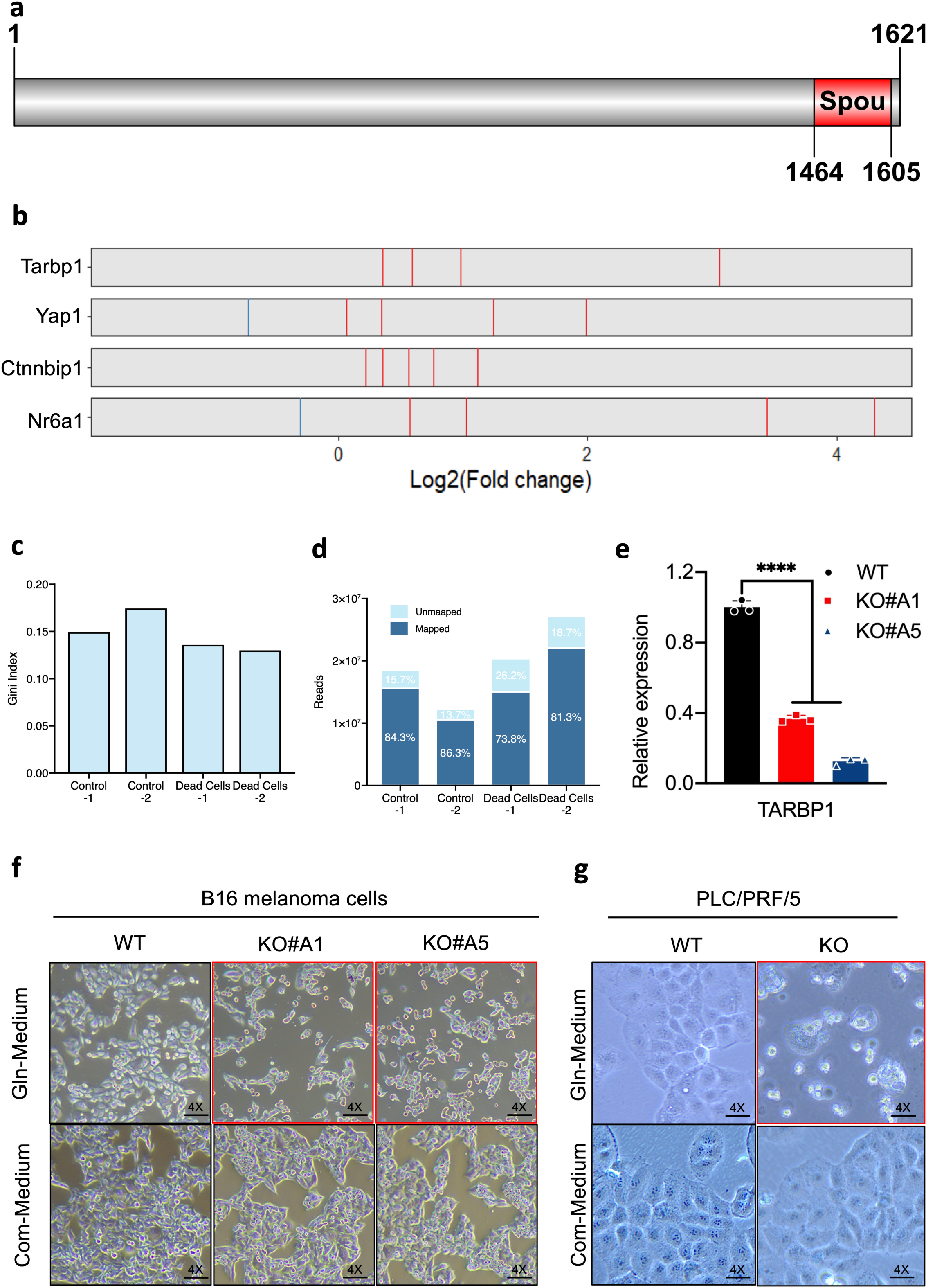
CRISPRi screens for essential genes in glutamine metabolism. a, A scheme showing functional domains of human TARBP1. b, Down-regulated (blue) and up-regulated (red) sgRNAs in CRISPRi screens. c, d, The Gini Index and QC plots of CRISPR screening libraries. e, Knockout efficiency of TARBP1 in B16 melanoma cells was testified by Real-time quantitative PCR analysis (No TARBP1 antibody working in WB with mouse samples). ****P<0.0001. Student’s t-test. Data represented as mean ± SD (n=3). f,g, Representative cell pictures of the WT and TARBP1 KO in B16 melanoma cells (f) and PLC/PRF/5 HCC cells (g) culturing in Glutamine-basic medium or Complete medium for 48 hours.

**Extended Data Fig. 2:**
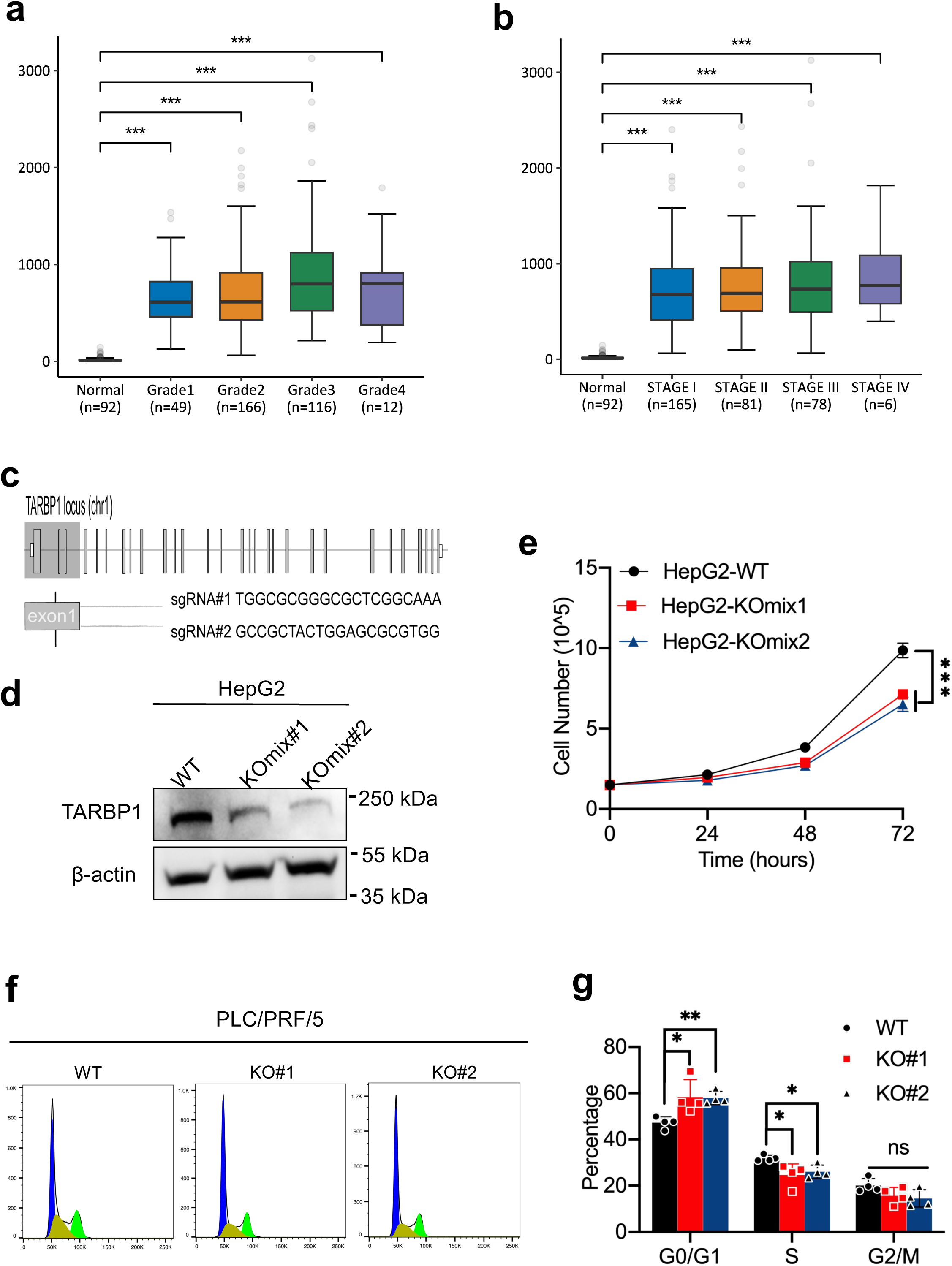
Deletion of TARBP1 impairs HCC progression. a, b, Correlations of TARBP1 levels with the tumor grade (a) and tumor stage (b) of HCC. c, Schematic diagram of sgRNA targeting human TARBP1 locus. d, Western blot analysis confirming TARBP1 KO pool in HepG2 HCC cell line with TARBP1 antibody. β-actin was used as a loading control. e, Cell proliferation analysis of WT and TARBP1 KO pool in HepG2 cells. f, g, Representative images (g) and quantification analysis (h) of cell cycle analysis of PLC/PRF/5 cells comparing TARBP1 KO versus WT control. *P<0.05, **P<0.01, ***P<0.001, ns: not significant. Student’s t-test. a,b, Data were presented as mean ± SEM. c, f, Data represented as mean ± SD (n=3). h, Data represented as mean ± SD (n=4).

**Extended Data Fig. 3:**
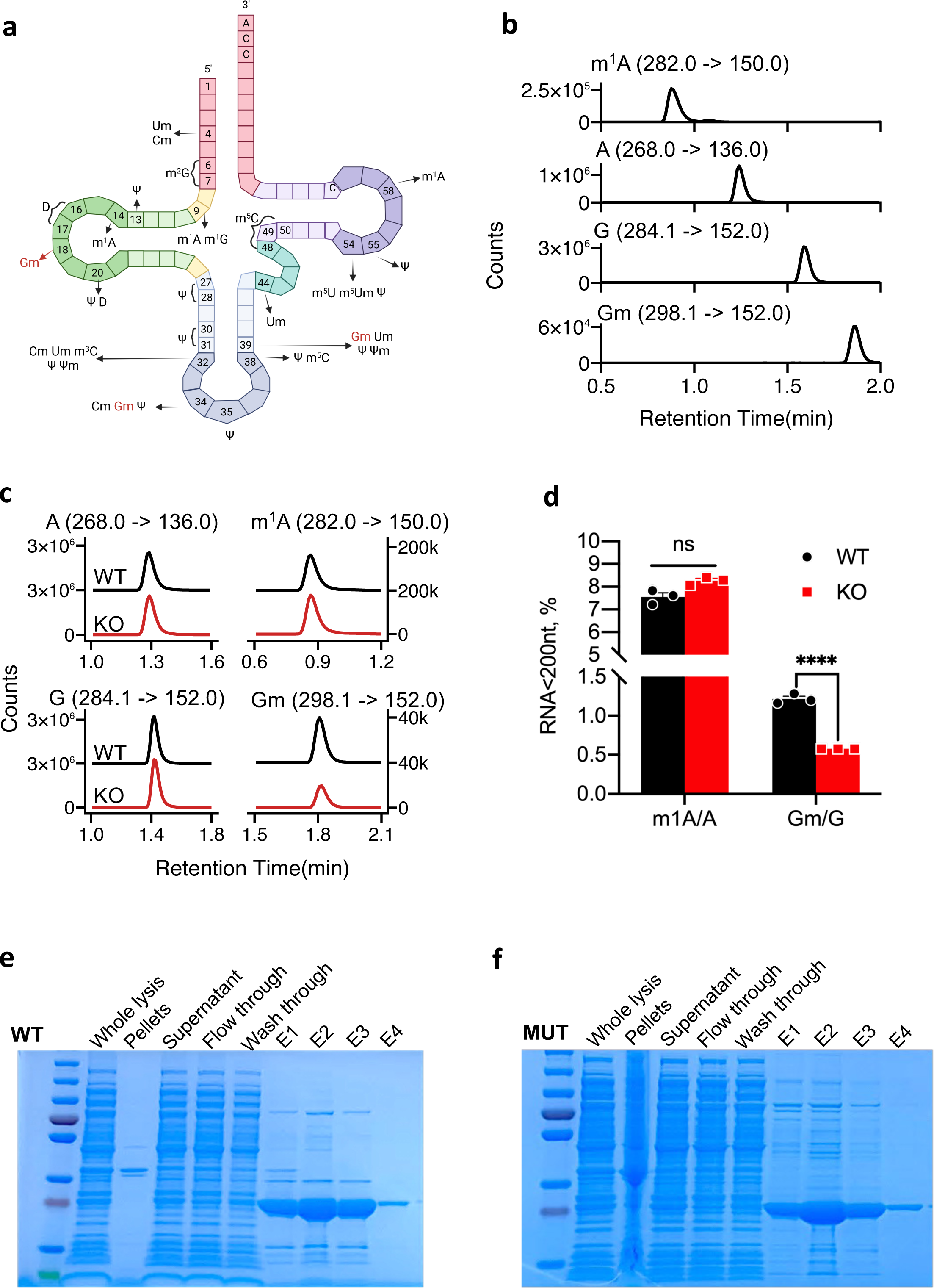
TARBP1 specifically methylates tRNA^Gln^ ^(TTG/CTG)^ and tRNA^ser^ ^(TGA/GCT)^ in vivo and in vitro. a, Schematic representation of a tRNA molecule and examples of RNA modifications. 2’-O-methylation at position 18 is highlighted in red. b, Retention time and nucleoside to base ion mass transitions of standard samples tested by LC-MS/MS. c, d, LC-MS/MS analysis of Gm/G levels in small RNA purified from the WT and TARBP1 KO cells (c). N=3 independent experiments (d). ****P<0.0001, ns: not significant. Student’s t-test. Data were presented as mean ± SEM. e, f, SDS-PAGE gels maps of recombinant protein purification, including truncated WT (e) and Mut TARBP1 protein (f).

**Extended Data Fig. 4:**
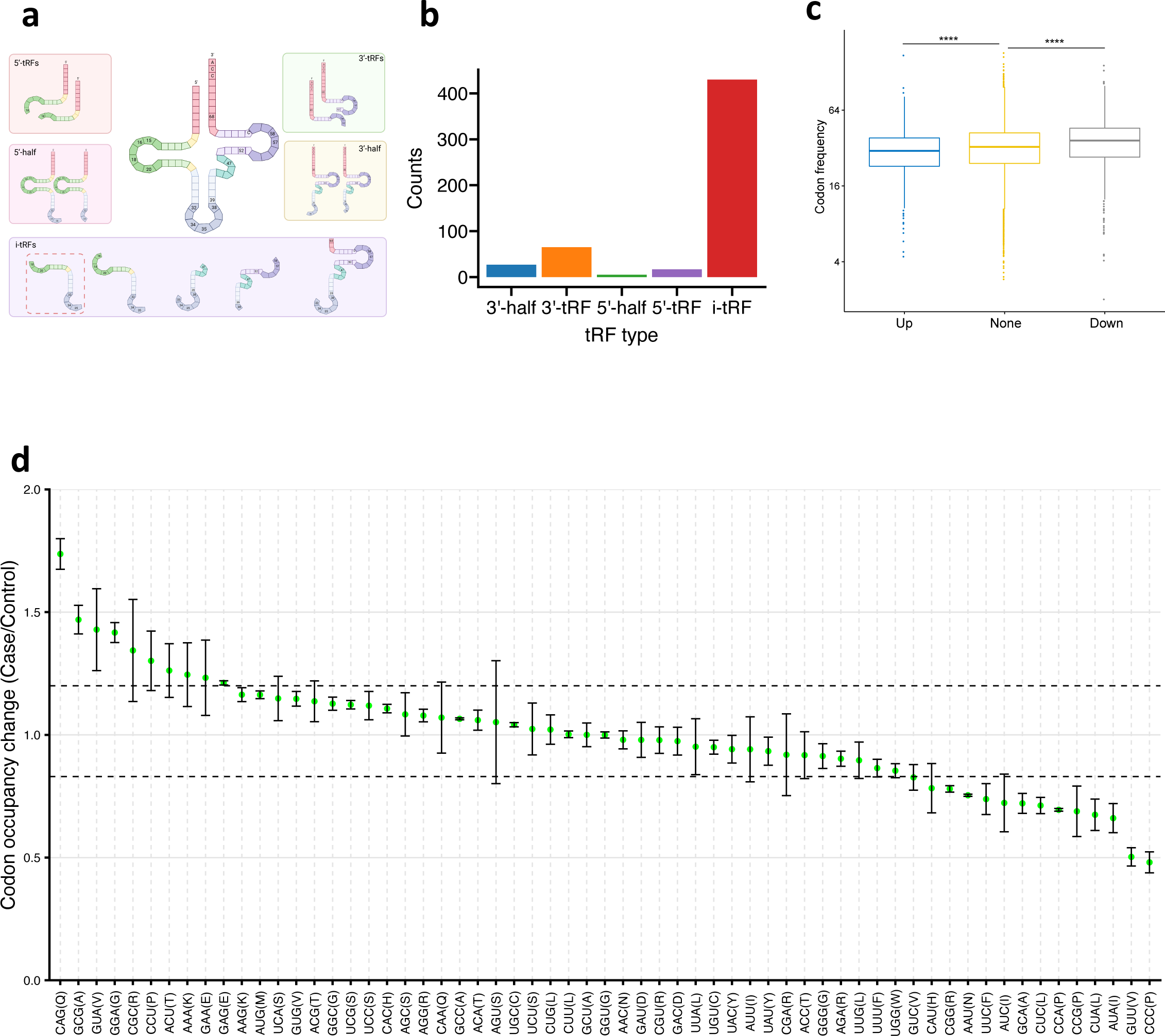
TARBP1 regulates gene expression via increasing the stability of Gm18 modified tRNAs. a, Schematic diagram of the five structural categories of tRNA-derived RNA fragments (tRFs). b, tRF type analysis of the increased tRFs upon loss of TARBP1. c, Overall CAG codon occupancy among the proteins with decreased translation efficiency (TE, down), unchanged TE (None), and increased TE (UP) in TARBP1 KO cells compared to WT cells. d, Codon occupancy at P sites in TARBP1 KO cells compared to WT cells.

**Extended Data Fig. 5:**
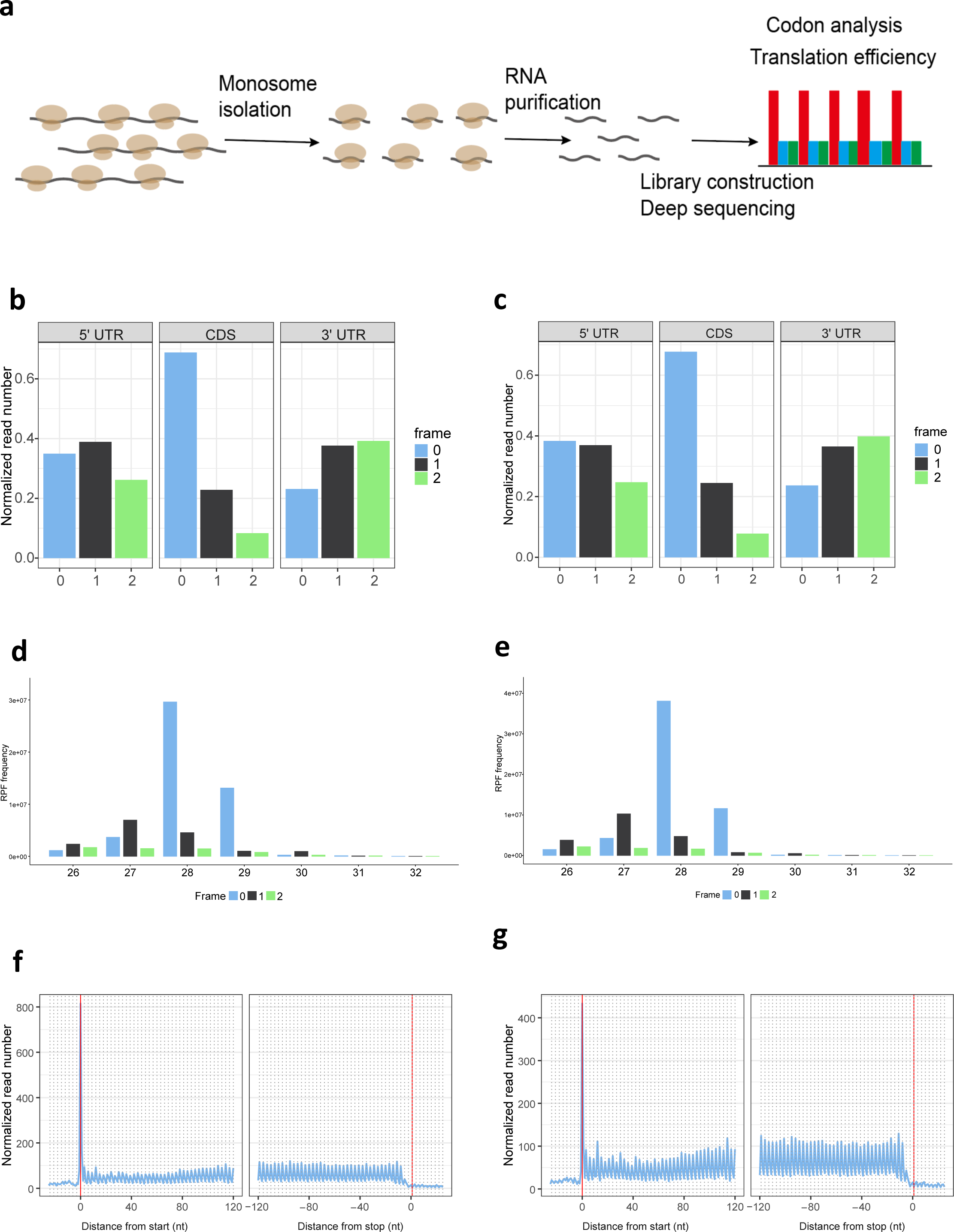
Ribo-seq measures global translation. a, Schematic diagram illustrating the principle of Ribo-seq. b-g, Characteristics of Riboseq data in control and TARBP1 KO cells. Ribosome protected fragment (RPF) length distribution of TARBP1 KO cells (b) and WT cells (c). Normalized A-site signals of RPFs for different coding frames among the 5’-UTR, coding sequence (CDS) and 3’-UTR of TARBP1 KO cells (d) and WT cells (e). Normalized A-site signals of RPFs near the translation start sites (TSS) and translation end sites (TES) of TARBP1 KO cells (f) and WT cells (g).

**Extended Data Fig. 6:**
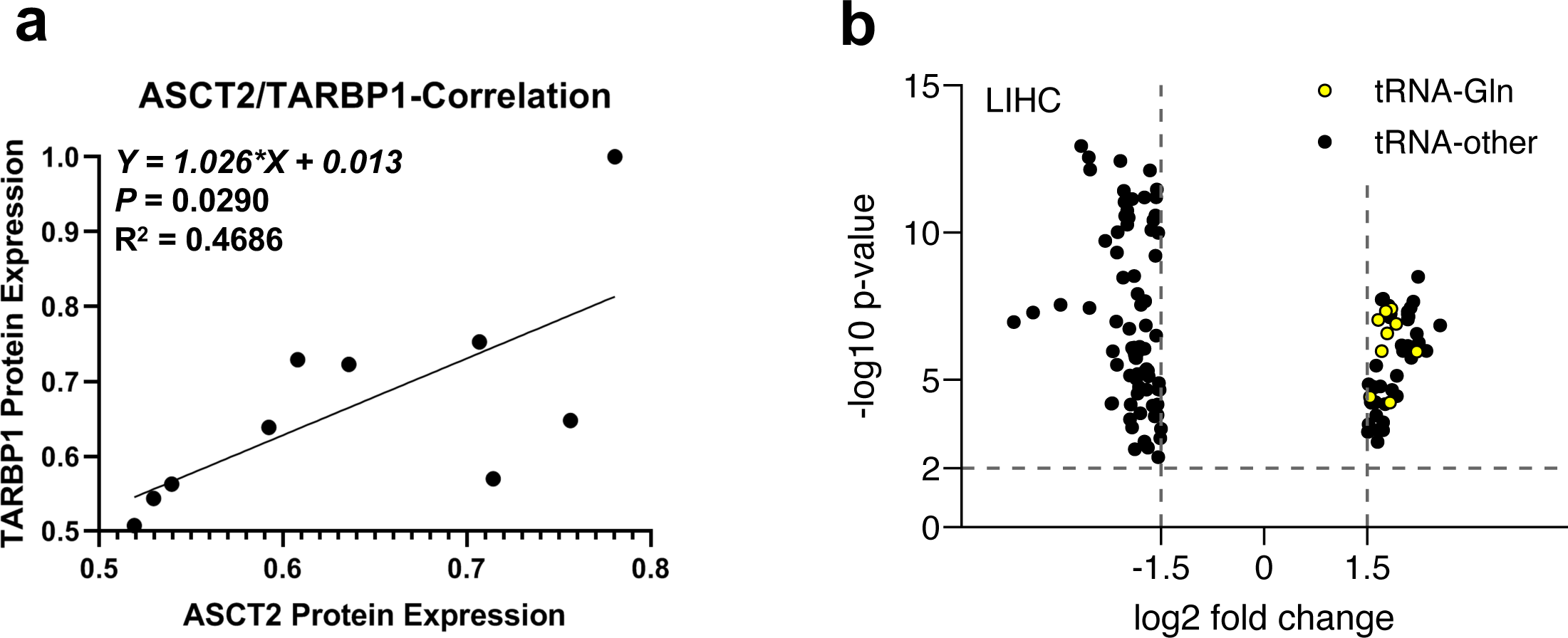
TARBP1 expression affects HCC progression. a, Correlations of TARBP1 expression with ASCT2 expression. b, tRNA levels in hepatobiliary cancers. Glutaminyl transfer RNA (tRNA-Gln) is highlighted in yellow.

